# Restoring Aquaporin-4 alleviates molecular and motor phenotypes in zQ175 mouse model of Huntington’s disease

**DOI:** 10.64898/2026.09.20.753038

**Authors:** Hongshuai Liu, Zichen Ding, Yadi Wang, Helen Moniz, Yuguo Li, Yuxiao Ouyang, Chang Liu, John Anderson, Jun Hua, Jiadi Xu, Wenzhen Duan

## Abstract

Huntington’s disease (HD) is characterized by progressive accumulation of mutant huntingtin (mHTT), neurodegeneration and motor dysfunction. Increasing evidence implicates impaired glymphatic function and loss of perivascular aquaporin-4 (AQP4) polarization in HD, suggesting that defective brain waste clearance may contribute to disease pathogenesis. Whether restoration of AQP4-mediated glymphatic transport can ameliorate HD pathology remains unknown.

We tested whether genetic restoration of AQP4 improves glymphatic function and HD-associated phenotypes in heterozygous zQ175 knock-in mice. At 2 months of age, premanifest zQ175 mice received a single retro-orbital injection of BBB-penetrant AAV9-PHP.eB encoding the AQP4 (M1 isoform). Glymphatic transport was assessed by intracisternal injection of fluorescent BSA-647 and quantification of parenchymal influx and drainage to the deep cervical lymph nodes. Perivascular AQP4 localization, mHTT aggregation, gliosis, DARPP32 immunoreactivity, CSF neurofilament light chain (NfL) and total Tau, structural brain volumes by 11.7-T MRI, and motor behavior were assessed longitudinally through 12 months of age.

AAV9-PHP.eB-AQP4 produced widespread brain expression and restored perivascular AQP4 localization in the striatum and cortex of zQ175 mice. AQP4 overexpression significantly increased glymphatic tracer influx into brain parenchyma and BSA-647 drainage to the deep cervical lymph nodes. At 12 months, AQP4 restoration reduced CSF NfL and total Tau concentrations, mHTT aggregate number and mean aggregate size, and striatal Iba1-positive microglia and GFAP-positive astrocytes (P < 0.05). AQP4 overexpression also preserved striatal DARPP32 immunoreactivity and significantly attenuated striatal atrophy measured by MRI. Female zQ175 mice showed improved balance-beam performance and locomotor activity, while effects on motor performance in males did not reach statistical significance. AQP4 overexpression attenuated body-weight loss in both sexes. No detectable structural, behavioral or body-weight abnormalities were observed following sustained AQP4 overexpression in wild-type mice.

In summary, restoration of AQP4 enhances glymphatic–lymphatic transport and is associated with reduced proteopathic, neuroinflammatory and neurodegenerative phenotypes in zQ175 HD mice. These findings provide proof-of-concept that impaired glymphatic function is a modifiable component of HD pathology and support AQP4-mediated restoration of brain waste-clearance pathways as a potential disease-modifying therapeutic strategy.

## Introduction

Huntington’s disease (HD) is an autosomal dominant neurodegenerative disorder caused by an expanded CAG trinucleotide repeat in the *huntingtin* (*HTT*) gene ^1,2^. This mutation results in the production of toxic mutant huntingtin (mHTT) protein that is prone to misfolding and aggregation. Progressive accumulation of mHTT aggregates initially affects the striatum and later extends to the cortex and other brain regions, leading to neuronal degeneration and circuitry dysfunction. Clinically, HD is characterized by motor, cognitive, and psychiatric symptoms. While HTT-lowering therapies have shown promise, no FDA-approved treatment has yet been shown to modify disease progression ^3^. Therefore, identifying additional therapeutic strategies that target unique pathogenic mechanisms could enhance development of new disease-modifying treatments.

The brain possesses a unique waste clearance system known as the glymphatic system; it facilitates the convective exchange of cerebrospinal fluid (CSF) and interstitial fluid (ISF) along perivascular spaces to mediate the removal of interstitial solutes, including metabolic waste and neurotoxic macromolecules, into the peripheral lymphatic system ^4^. This brain-wide fluid transport pathway is highly dependent on the polarized expression of the water channel aquaporin-4 (AQP4) at the vascular-facing endfeet of astrocytes ^5,6^. The perivascular localization of AQP4 facilitates CSF influx into the brain parenchyma and supports clearance of ISF to the deep cervical lymph nodes ^7–10^. In Alzheimer’s disease and Parkinson’s disease, the loss of AQP4 polarization disrupts glymphatic flux ^8,11,12^. Impaired glymphatic transport may contribute to the accumulation of pathogenic proteins and consequently promote neurodegeneration by reducing the clearance of neurotoxic proteins and metabolites from the brain ^13,14^. We have previously discovered impaired glymphatic flow and reduced AQP4 perivascular polarization in premanifest zQ175 HD mice ^15^. Recent neuroimaging studies have identified altered CSF flow dynamics, implicating that disturbed glymphatic function correlates with disease severity in HD patients ^16–18^. These clinical findings support the idea that glymphatic dysfunction may serve as a potentially modifiable pathological mechanism in HD ^17–19^. Whether targeting this pathway can exert a therapeutic effect in HD has not been explored.

In this study, we administered an AAV9-PHP.eB vector encoding GFP-tagged AQP4 (M1 isoform) by retro-orbital injection to zQ175 HD mice at 2 months of age. Through a combination of analysis of glymphatic function by tracking the fluorescent CSF tracer movement and HD-relevant pathological and behavioral assessment following AQP4 overexpression, we found that AQP4 overexpression restored perivascular AQP4 localization in zQ175 HD mice, improved glymphatic transport, and attenuated HD-associated neuropathology and motor deficits. At the same time, it had no detectable adverse effects in wild-type control mice. These findings support AQP4-mediated restoration of glymphatic transport as a potential disease-modifying strategy for HD.

## Materials and methods

### Animals

Heterozygous zQ175 knock-in mice (The Jackson Laboratory, Bar Harbor, ME, USA) and their age-matched wild-type (WT) littermates were used in this study. Genotyping and CAG repeat length determination were performed by PCR analysis of tail biopsy samples. The zQ175 mice used in this study carried approximately 220 ± 5 CAG repeats. Mice were housed under standard laboratory conditions on a 12-h light/dark cycle, with ad libitum access to food and water.

All animal procedures were approved by the Johns Hopkins University Institutional Animal Care and Use Committee (IACUC) and conducted in accordance with the National Institutes of Health Guide for the Care and Use of Laboratory Animals. Behavioral testing and MRI experiments were conducted during the dark phase of the light/dark cycle. Mice were anesthetized with isoflurane during MRI acquisition, as described below.

Mice were randomly assigned to the experimental groups, and littermate controls were used for genotype comparisons whenever possible. Animals from multiple litters were included in each experimental group to minimize litter-specific effects. Investigators blinded to genotype and treatment performed data collection and quantitative analyses using coded animal identifiers. Outliers were excluded only in rare instances in which technical artifacts or unexpected animal death precluded reliable data acquisition.

### AAV vector production and administration

The AAV vector encoding mouse AQP4-M1 fused to GFP under the control of the GFAP promoter was packaged in the AAV9-PHP.eB capsid. Control mice received an equivalent dose of AAV expressing GFP alone. Mice at 8 weeks of age received a single retro-orbital injection of AAV at a dose of 1.5 × 10¹¹ vg per mouse in a total volume of 100 μL. For early validation of transgene expression, mice were analyzed 4 weeks after AAV administration. For long-term studies, glymphatic transport, MRI, behavioral testing, and biochemical analyses were performed at the indicated ages.

### Immunofluorescence staining and imaging

Mice were anesthetized with isoflurane and transcardially perfused with phosphate-buffered saline (PBS), followed by 4% paraformaldehyde (PFA). Brains were collected, post-fixed overnight in 4% PFA, and subsequently cryoprotected in 30% sucrose for 24 h. Coronal brain sections (40 μm) were prepared using a microtome.

Free-floating coronal sections were washed three times with PBS for 10 min per wash and permeabilized with 0.3% Triton X-100 in PBS for 5 min. Sections were blocked for 1 h at room temperature in PBS containing 5% donkey serum, 5% goat serum, and 0.1% Triton X-100, then incubated overnight at 4°C with primary antibodies against AQP4 (249-323; 1:100; Alomone Labs), CD31 (AF3628; 1:100; Bio-Rad), GFAP (13-0300; 1:100; Invitrogen, Thermo Fisher Scientific), EM48 (MAB5374; 1:100; Sigma-Aldrich), Iba1 (019-19741; 1:100; FUJIFILM Wako), and DARPP-32 (2306; 1:100; CST). The following day, sections were washed three times with PBS and incubated with appropriate fluorescent secondary antibodies for 1 h at room temperature. Sections were counterstained with DAPI, mounted on Superfrost slides, and coverslipped with SlowFade Gold Antifade Mountant. All steps, including and after secondary antibody incubation, were performed protected from light.

Fluorescence images were acquired using a Zeiss LSM 700 confocal microscope equipped with an Axio Observer platform. Identical acquisition settings were used for all experimental groups. Images were acquired at a resolution of 1,024 × 1,024 pixels and an in-plane pixel size of 0.2 μm per pixel. Depending on the staining panel, images were acquired in the DAPI, 488-nm, 555-nm, and 647-nm channels.

### Image analysis and quantification

Samples were assigned unique identifiers, and investigators performing image acquisition and analysis were blinded to genotype and treatment groups. Group identities were revealed only after image quantification was complete. For each animal, three microscopic fields from three anatomically matched coronal sections were acquired and analyzed, and the mean value was used as a single biological replicate.

### AQP4 perivascular localization

Perivascular localization of AQP4 was quantified by measuring its colocalization with the vascular marker CD31 using ZEN 3.4 Blue Edition (Carl Zeiss). Identical acquisition settings and threshold parameters were used for all experimental groups. Colocalization was quantified within a 160 × 160 μm field of view acquired with a 20× objective and reported as AQP4–CD31 colocalization density normalized to CD31-positive vascular area. The mean value from three sections per mouse was used for statistical analysis.

### Quantification of neuroinflammatory and striatal markers

Iba1-positive cells and GFAP-positive astrocytes were quantified in anatomically matched striatal sections using identical acquisition settings and thresholding criteria across groups. Cell counts were normalized to ROI area and reported as cells/mm². DARPP32 immunoreactivity was quantified as mean fluorescence intensity within the striatal ROI and normalized to WT-Con mice. For each mouse, three coronal sections were selected and analyzed, and the mean value was treated as a single biological replicate.

### mHTT aggregate analysis

Mutant huntingtin (mHTT) aggregates were quantified using a custom Python-based image analysis pipeline. Fluorescence images contained three channels, in which the AF555 channel represented mHTT immunoreactivity, and the AF647 channel represented the AQP4 signal. Since the AF555 channel exhibited partial spectral overlap with the AF647 channel representing the AQP4 signal, an exclusion mask was generated from the AF647 channel to remove any AQP4-associated fluorescence signals before aggregate segmentation. A fixed exclusion threshold was applied to the AF647 channel: pixels with AF647 fluorescence intensities exceeding 4669, corresponding to the top 2% of fluorescence intensities across all images, were excluded from the AF555 channel without morphological dilation.

After signal exclusion, mHTT aggregates were segmented in the AF555 channel using fixed hysteresis thresholds (low threshold = 2219.91; high threshold = 3934.00), which were calculated from the image datasets and applied uniformly to all images. Pixels below the low threshold were classified as background, whereas those above the high threshold were classified as aggregate-positive. Spatially connected pixels were identified as individual aggregates by connected-component labeling. Quality-control images were generated to verify the performance of the exclusion mask and aggregate segmentation.

Three quantitative parameters were extracted for each image: (i) aggregate number, (ii) total aggregate area, and (iii) mean aggregate area. Aggregate areas were converted from pixels to μm² using spatial calibration metadata extracted from the original CZI image files.

### Intracisternal tracer injection

To track CSF movement and glymphatic transport in the CNS, mice were anesthetized with isoflurane and positioned in a stereotaxic frame. The cisterna magna was surgically exposed, and a 30-gauge needle was inserted into the cisterna magna. Alexa Fluor 647-conjugated bovine serum albumin (BSA-647; Invitrogen, Thermo Fisher Scientific; 66 kDa) was dissolved in artificial cerebrospinal fluid (aCSF) at a concentration of 0.5% (w/v). Each mouse received 10 μL of BSA-647 infused at 2 μL/min over 5 min using a syringe pump (Harvard Apparatus). All injections were performed by the same operator.

One hour after intracisternal injection, mice were transcardially perfused. Brains were collected and coronally sectioned at 60 μm using a Leica cryostat. Sections were mounted on glass slides and imaged ex vivo using a Zeiss fluorescence microscope. Tiled images were acquired to reconstruct the entire coronal brain section under identical imaging settings across experimental groups.

Tracer influx was quantified using Fiji (ImageJ) by investigators blinded to experimental groups, as previously described^4^. The brain parenchyma in each coronal section was manually delineated as the region of interest (ROI), and thresholded BSA-647-positive area coverage within the ROI was quantified using consistent thresholding parameters across all samples. Four anatomically matched coronal sections were analyzed per mouse, and the mean value was used as the representative value for each mouse.

To assess downstream lymphatic drainage, deep cervical lymph nodes (dCLNs) were collected 1 h after intracisternal BSA-647 injection. dCLNs were imaged under identical acquisition settings across experimental groups, and BSA-647-positive area coverage within the lymph node ROI was quantified using Fiji with identical thresholding parameters across experimental groups.

### Structural MRI acquisition and image analysis

In vivo structural MRI was performed on a horizontal-bore 11.7 T Bruker BioSpec system equipped with a 72-mm quadrature transmit coil and a four-element phased-array receive coil. Structural brain images were acquired at a spatial resolution of 0.1 × 0.1 × 0.2 mm³.

Images were analyzed using an established pipeline developed previously in our laboratory. Skull-stripped images were rigidly aligned and processed using Landmarker software. Intensity-normalized images were submitted for large-deformation diffeomorphic metric mapping. Regional brain volumes were quantified from the Jacobian determinants of the transformations generated by LDDMM. All MRI data were analyzed by the same investigator to minimize variability.

### Animal preparation and physiological monitoring

Each mouse was anesthetized with 2% isoflurane and maintained with 1%–1.5% isoflurane throughout MRI acquisition. Each mouse was positioned on a water-heated bed, and the head was immobilized using a bite bar and head holder. Throughout the experiment, the respiratory rate of each mouse was continuously monitored using a physiological monitoring system (SA Instruments) and maintained at 40– 60 breaths/min.

### CSF collection and biomarker analysis

For each mouse, the CSF was collected from the cisterna magna under isoflurane anesthesia at the end of the experiment. The CSF samples were immediately placed on ice after collection, briefly centrifuged to remove cellular debris, and stored at −80°C until analysis. Visibly blood-contaminated samples were excluded from analysis.

CSF concentrations of neurofilament light chain and total Tau were measured using the Meso Scale Discovery S-PLEX Neurology Panel 1 kit according to the manufacturer’s instructions. Samples were analyzed in duplicate, and biomarker concentrations were calculated from standard curves generated on the same assay plate. All measurements were performed by investigators blinded to the genotype and treatment group.

### Behavioral tests

Balance beam. The balance beam test was performed to assess motor coordination in each mouse. Motor coordination and balance were evaluated using an 80-cm-long, 5-mm-wide square balance beam elevated 50 cm above the surface. A bright light was positioned at the starting platform, and a darkened enclosed escape box (12 × 12 × 12 cm; 1,728 cm³) was placed at the opposite end of the beam. Mice underwent two training trials on the beam one day before testing. During the balance beam test, the time for each mouse to traverse the entire length of the beam was recorded. Mice that fell from the beam during testing were assigned a maximum cutoff latency of 125 s.

Locomotor activity. Spontaneous locomotor activity was assessed in a 40 × 40 × 40 cm acrylic open-field arena during the dark phase under no visible illumination. Mice were tested individually, placed in the center of the arena, and recorded for 60 min at 7 frames/s. Testing order was randomized, and the arena was cleaned between animals. Behavioral tracking and analysis were performed by investigators blinded to group assignments.

Videos were analyzed using ToxTrac 2026. Locomotor activity was quantified during a prespecified 28-min window from 20 to 48 min after placement to allow habituation. Total distance traveled was used as the primary measure of spontaneous locomotor activity, and tracking quality was verified by visual inspection.

### Statistics

Data are presented as mean ± standard error of the mean (SEM) unless otherwise indicated. Statistical analyses were performed using GraphPad Prism 10 (GraphPad Software, San Diego, CA, USA). Comparisons between two groups were performed using unpaired two-tailed Student’s t-tests. Comparisons among multiple groups were performed using one-way or two-way analysis of variance (ANOVA), as appropriate, followed by Bonferroni’s post hoc test for multiple comparisons. Behavioral data were analyzed separately by sex. Longitudinal body weight data were analyzed separately by sex using two-way repeated-measures ANOVA. P < 0.05 was considered statistically significant. Individual data points are shown in the corresponding figures.

## Results

### Systemic delivery of AAV-PHP.eB-AQP4 facilitates brain-wide expression and perivascular polarization in HD mice

To evaluate the effects of restoring AQP4 levels on HD progression, we administered AAV9-PHP.eB-AQP4 (M1 isoform) fused with GFP via retro-orbital injection to 2-month-old (premanifest) zQ175 mice. Glymphatic function was assessed at 7 months of age by tracking intracisternal-injected CSF fluorescent tracer BSA-647 movement to quantify glymphatic influx and efflux to the deep cervical lymph nodes (dCLNs). Structural MRI for brain volumetric assessment was performed at 10 months of age. Motor behavioral, biochemical, and histological assays were performed at 12 months of age in zQ175 mice. The experimental timeline is illustrated in **Fig. 1A**.

**Figure 1.**
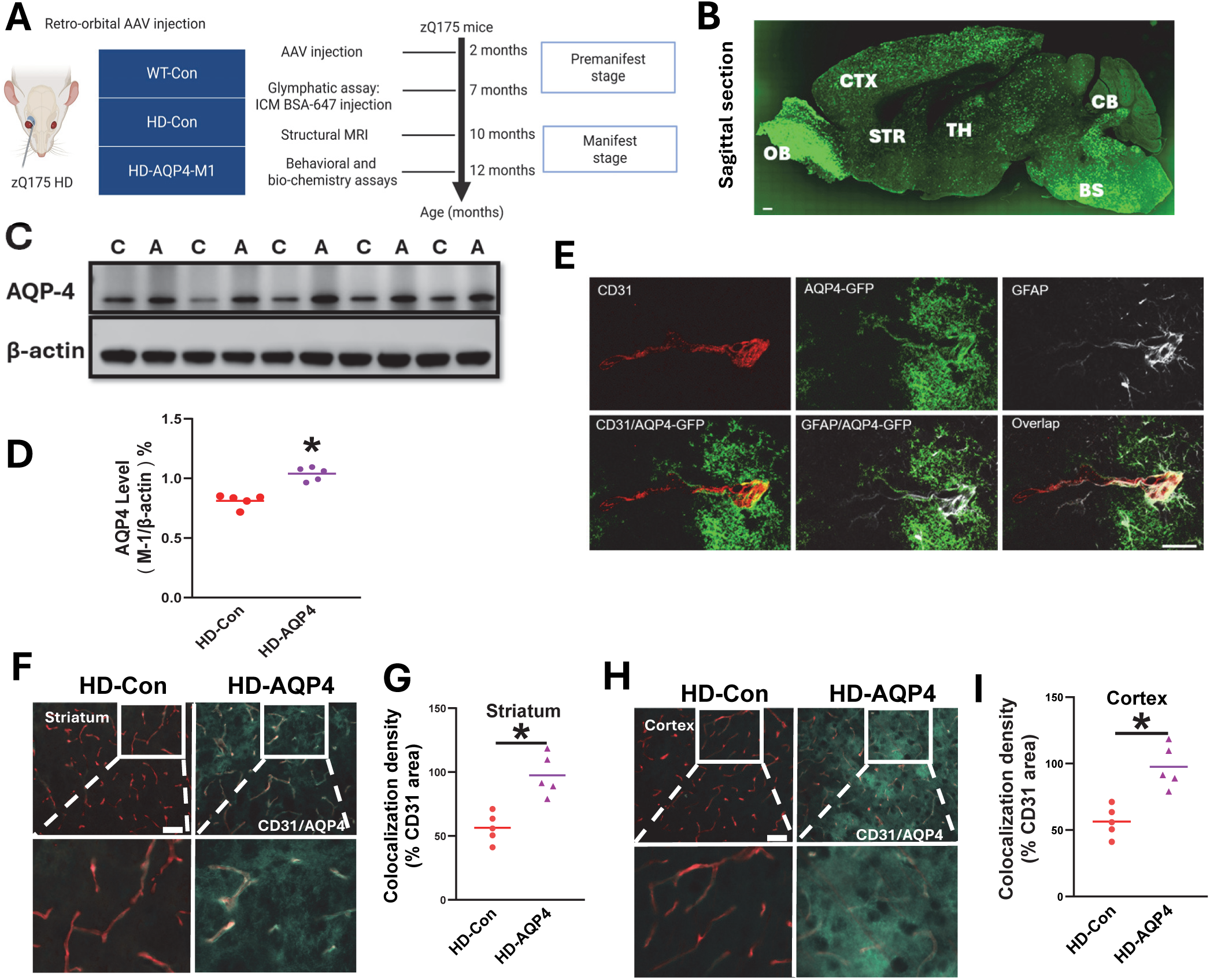
Characterization of AAV9-PHP.eB-mediated AQP4 expression and perivascular localization in 3-month-old zQ175 HD mice. (**A**) Illustration of experimental design. (**B**) A representative sagittal brain section shows widespread transgene AQP4 (GFP fluorescence) following systemic delivery of AAV-PHP.eB-AQP4. Transgene expression was detected throughout the brain, including the olfactory bulb (OB), cortex (CTX), striatum (STR), thalamus (TH), brainstem (BS), and cerebellum (CB). (**C**) Western blot analysis of AQP4 in zQ175 HD mice receiving control virus (C) or AQP4 virus (A). (**D**) Quantification of AQP4 protein levels normalized to β-actin. (**E**) Representative confocal images showing CD31 immunosignal (red), AQP4 transgene (GFP, green), and GFAP-positive astrocytes (white). Note that transgene AQP4 is localized at the endfeet of astrocytes. (**F**) Representative confocal images of CD31 (red) and AQP4 (green) in the striatum of zQ175 HD mice (HD) receiving control or AQP4 AAVs. (**G**) Quantification of colocalization of AQP4 and CD31 signals in the striatum. (**H**) Representative confocal images of CD31 (red) and AQP4 (green) in the cortex of zQ175 HD mice (HD) receiving control or AQP4 AAVs. (**I**) Quantification of colocalization of AQP4 and CD31 signals in the cortex. Individual data points are shown in all graphs, and statistical significance was determined using an unpaired Student’s t-test. \**P* < 0.05.

We first determined whether systemic delivery (i.v.) of AAV-PHP.eB-AQP4 produced widespread expression in the zQ175 mouse brain. One month after AAV delivery, we observed robust GFP fluorescence throughout the brain, including the olfactory bulb, cortex, striatum, thalamus, brainstem, and cerebellum (**Fig. 1B**), indicating broad expression of exogenous AQP4 following AAV-PHP.eB administration. Western blot analysis further confirmed increased AQP4 protein levels in HD mice injected with AAVs carrying the AQP4 construct (**Fig. 1C, D**).

High-magnification confocal microscopy revealed that the overexpressed AQP4 (GFP fluorescence) is localized alongside astrocytic endfeet. Specifically, triple immunofluorescence staining for CD31 (red, vascular endothelial cell marker), GFP (green, exogenous AQP4), and GFAP (white, astrocyte marker) illustrated localization of the overexpressed AQP4, including the part localized to the perivascular astrocytic endfeet (**Fig. 1E**). Immunohistochemical analysis of AQP4 further demonstrated increased perivascular-associated AQP4 staining in the striatum (**Fig. 1F, G**) and cortex (**Fig. 1H, I)** following AAV-PHP.eB-AQP4 administration in zQ175 HD mice. These data confirm that systemic delivery of AAV9-PHP.eB-AQP4 promotes broad overexpression of AQP4 in the zQ175 mouse brain and facilitates its perivascular polarization in HD condition.

### AQP4 overexpression improves glymphatic transport and reduces accumulation of neuronal injury markers in the CSF of HD mice

To determine whether the restored AQP4 levels and its perivascular localization translated into functional improvements in glymphatic transport, we performed an assay to evaluate glymphatic flow by injecting a fluorescent tracer, BSA-647, into the cisterna magna (ICM) at 5 months after AAV delivery. Compared with wild-type (WT) mice, HD mice exhibited reduced CSF tracer penetration along the perivascular space into the brain parenchyma (**Fig. 2A, B**), indicating impaired glymphatic influx in the HD condition. In contrast, HD mice administered with AAV-AQP4 exhibited a significant rescue in parenchymal tracer distribution, indicating improved glymphatic tracer influx following AQP4 overexpression in zQ175 HD mice (**Fig. 2A, B**).

**Figure 2.**
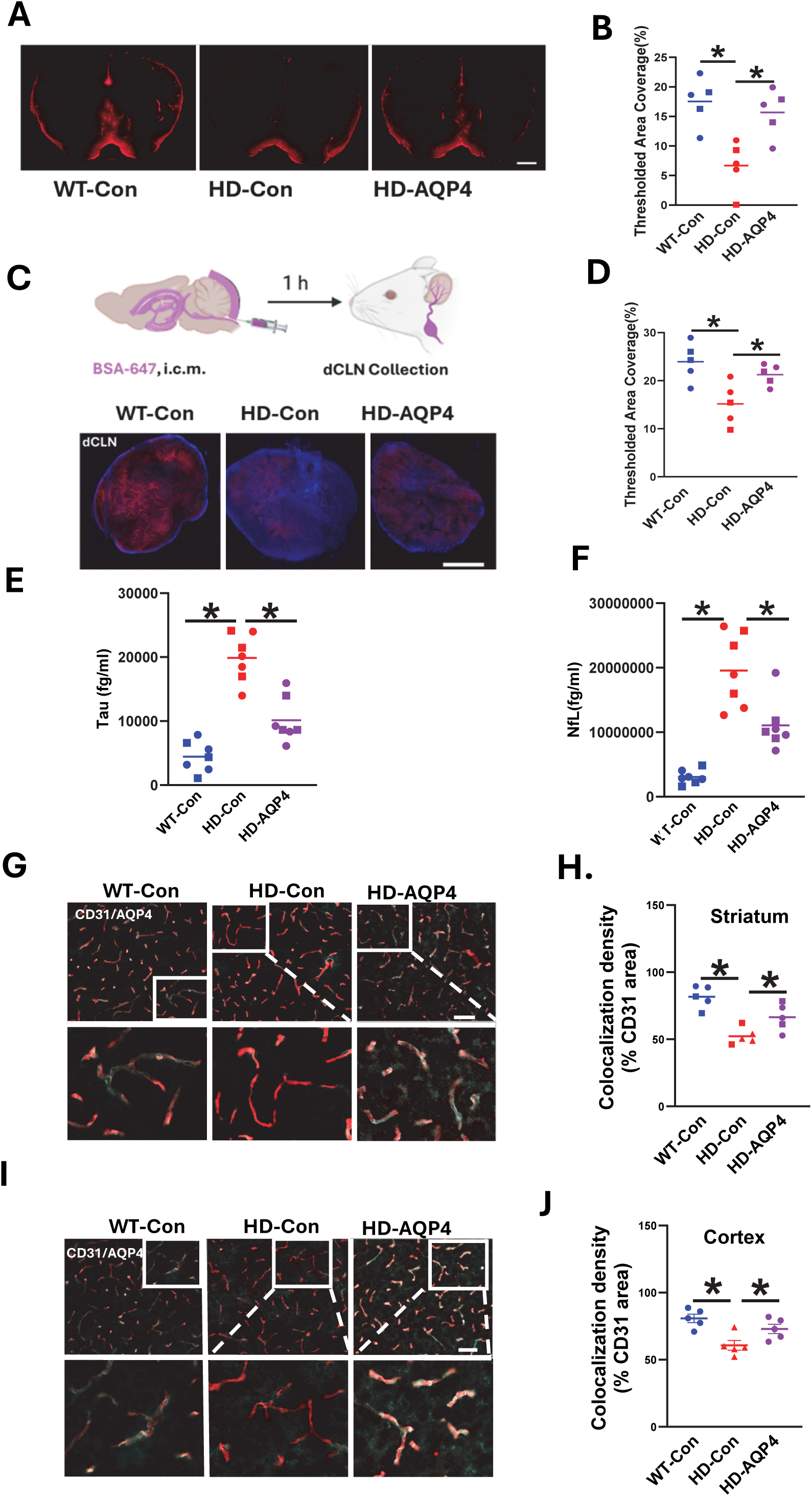
AQP4 overexpression improves glymphatic–lymphatic tracer transport and reduces CSF neuronal injury markers in HD mice. **(A)** Representative images showing distribution of intracisternally injected fluorescent tracer BSA-647 in indicated groups. Scale bar = 1 cm. **(B)** Quantification of thresholded BSA-647 area coverage in the brain parenchyma of 7-month-old mice. **(C)** Upper panel: experimental schematic showing intracisternal injection of BSA-647 followed by collection of deep cervical lymph nodes (dCLNs). Bottom panel: Representative fluorescence images of dCLNs from indicated groups. Scale bar = 350 µm. **(D)** Quantification of thresholded BSA-647 area coverage in dCLNs from 7-month-old mice. **(E, F)** Quantification of total Tau (E) and neurofilament light chain (NfL, F) concentrations in the CSF of 12-month-old mice from indicated groups. **(G)** Representative confocal images of the striatum showing CD31-positive blood vessels (red) and AQP4 immunoreactivity (cyan). Insets show higher-magnification views of the boxed regions illustrating perivascular AQP4 localization. Scale bar = 50 µm. **(H)** Quantification of AQP4 perivascular localization (colocalization with CD31) in the striatum of 12-month-old mice from indicated groups. **(I)** Representative confocal images of the cortex showing CD31 (red) and AQP4 (cyan) immunoreactivity. Insets show higher-magnification views of the boxed regions illustrating perivascular AQP4 localization. **(J)** Quantification of AQP4 perivascular localization in the cortex of 12-month-old mice from indicated groups. In all graphs, individual data points are shown, statistical significance was determined using one-way *ANOVA* followed by Bonferroni’s test. \**P* < 0.05.

Because dCLNs serve as a major downstream drainage site for cranial glymphatic effluent, dCLNs were collected 1 hour after ICM BSA-647 injection in 7-month-old mice, corresponding to 5 months after AAV-AQP4 delivery. BSA-647 accumulation in the dCLNs was markedly reduced in HD mice compared with WT littermate controls. In contrast, AQP4 overexpression significantly increased BSA-647 fluorescence in the dCLNs of zQ175 HD mice (**Fig. 2C, D**), indicating improved downstream glymphatic efflux and drainage to peripheral lymph nodes by AQP4 overexpression in HD mice.

We next examined whether AQP4 overexpression had an impact on the accumulation of neuroaxonal injury markers in the CSF. Concentrations of neurofilament light chain (NfL) ^20–23^ and total Tau ^24–27^ were measured at 12 months of age in the CSF of mice. zQ175 HD mice had significantly elevated CSF Tau and NfL levels compared with WT controls (**Fig. 2E, F**). AQP4 overexpression significantly reduced the concentration of both neuronal injury markers in the CSF (**Fig. 2E, F**), suggesting attenuation of neuroaxonal injury.

Because efficient glymphatic transport depends on the perivascular localization of AQP4, we assessed whether AQP4 overexpression maintained this perivascular localization by analyzing AQP4/CD31 colocalization in 12-month-old zQ175 mice, corresponding to 10 months after AAV delivery. Consistent with our previous findings ^15^, reduced AQP4 polarization was observed in the striatum (**Fig.2G, H**) and cerebral cortex (**Fig. 2I, J**) of 12-months-old zQ175 HD mice. AQP4 overexpression significantly restored perivascular AQP4 polarization, as indicated by increased colocalization of AQP4 and CD31 (**Fig.2G - J**). Together, these findings demonstrate that AQP4 overexpression enhances glymphatic influx and downstream drainage in HD mice and is associated with reduced CSF levels of neuronal injury-associated biomarkers.

### AQP4 overexpression reduces mHTT aggregation in HD mice

Having established that AQP4 overexpression improved glymphatic transport in HD mice, we next examined whether this intervention influenced mHTT accumulation in the brain. We performed immunofluorescent staining using the EM48 antibody in 12-month-old zQ175 mice, which recognizes the N-terminal region of mHTT and preferentially detects aggregated mHTT species. zQ175 HD mice exhibited a high density of mHTT aggregates throughout the striatum (**Fig. 3A**). Quantitative analysis showed a trend toward reduced total EM48-positive aggregate area (**Fig. 3B**), while both mHTT aggregate number (**Fig. 3C**) and average aggregate size (**Fig. 3D**) were significantly decreased in HD-AQP4 mice compared with HD controls. These findings suggest that AQP4 overexpression reduces striatal mHTT aggregate pathology in zQ175 HD mice.

**Figure 3.**
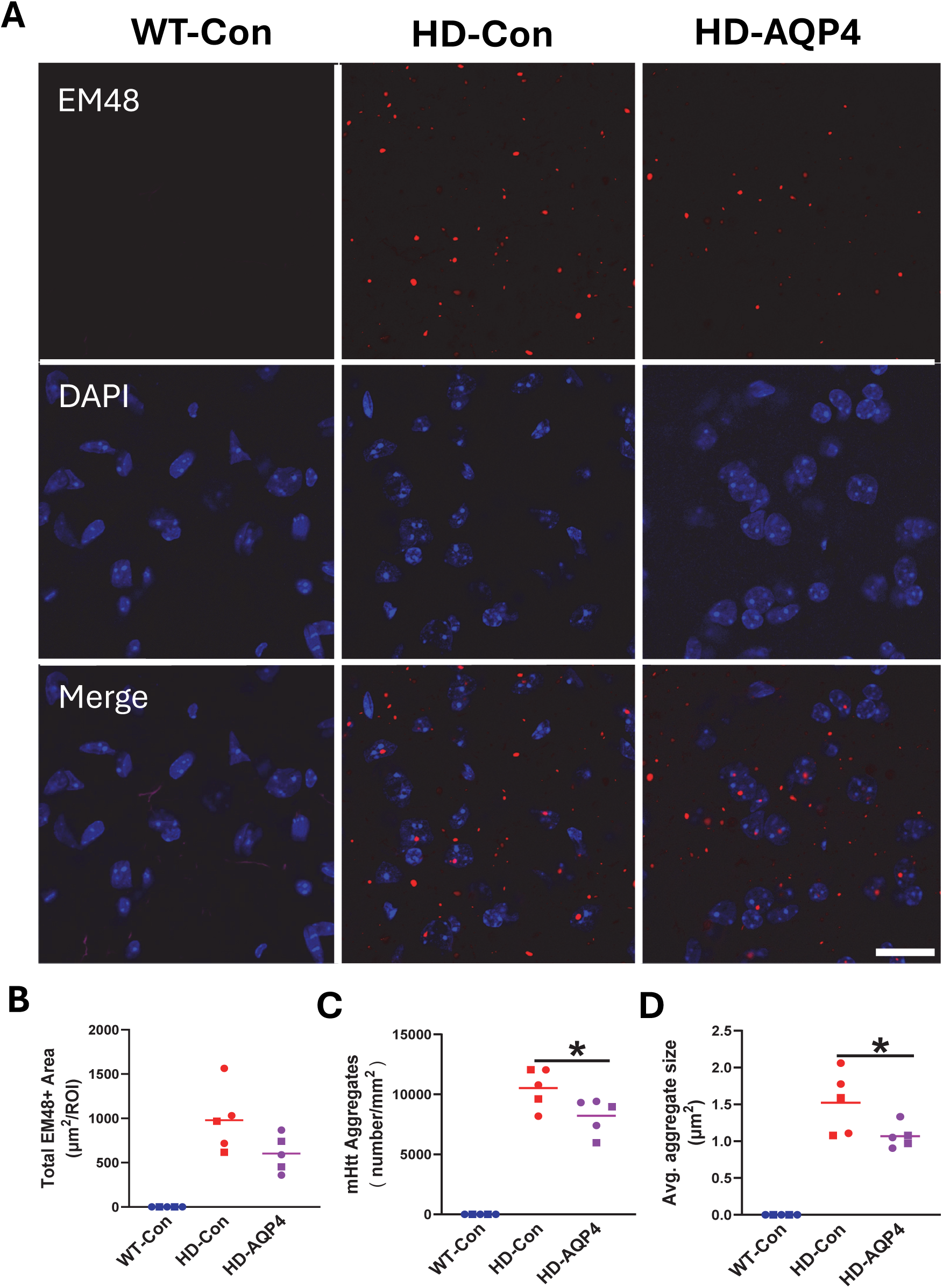
AQP4 overexpression reduces mutant huntingtin aggregation in zQ175 mice. **(A)** Representative images of EM48-positive immunofluorescence signals. Scale bar = 20 μm. **(B–D)** Quantification of mHTT aggregate burden in the striatum. AQP4 overexpression reduced mHTT aggregate burden in zQ175 mice, as shown by a decrease in total EM48-positive aggregate area (B) and significant reductions in aggregate number (C) and average aggregate size (D). Individual data points are shown, statistical significance was determined using one-way *ANOVA* followed by Bonferroni’s test. \**P* < 0.05.

### AQP4 overexpression reduces neuroinflammation and preserves striatal DARPP32 immunoreactivity in HD mice

Chronic neuroinflammation and reduced striatal DARPP32 expression are key pathological features of HD. To assess the effect of AQP4 overexpression on neuroinflammation, we performed immunostaining for microglia (Iba1) and activated astrocytes (GFAP). 12-month-old zQ175 HD mice showed increased Iba1-positive cell density (**Fig. 4A, B**) and GFAP-positive astrocyte density (**Fig. 4 C, D**) in the striatum, indicating increased gliosis and neuroinflammation, whereas AQP4 overexpression significantly reduced both measures (**Fig. 4A-D**).

**Figure 4.**
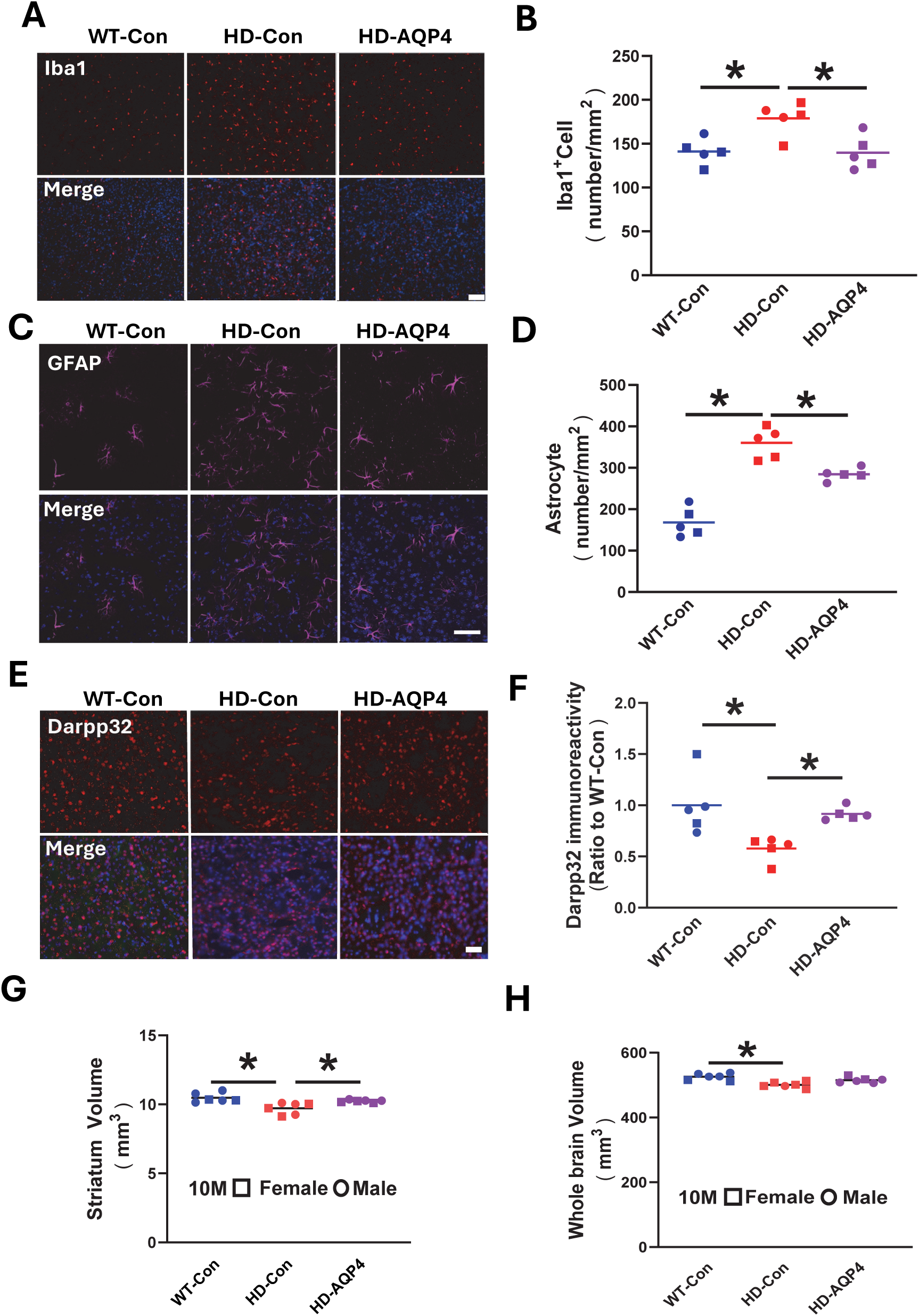
AQP4 overexpression attenuates neuroinflammation, preserves striatal DARPP32 immunoreactivity, and reduces striatal atrophy in zQ175 mice. **(A)** Representative Iba1 immunofluorescence images in the striatum. Scale bar = 50 µm. **(B)** Quantification of Iba1-positive cell density in the striatum. **(C)** Representative GFAP immunofluorescence images in the striatum. Scale bar = 20 µm. **(D)** Quantification of GFAP-positive astrocyte density in the striatum. **(E)** Representative striatal sections stained for DARPP32. Scale bar = 50 µm. **(F)** Quantification of DARPP32 immunoreactivity in the striatum. **(G–H)** Quantification of striatal volume (G) and whole-brain volume (H) by MRI scans. Individual data points are shown, statistical significance was determined using one-way ANOVA followed by Bonferroni’s test. \**P* < 0.05.

DARPP32 immunoreactivity, a marker of striatal medium spiny neuron-associated pathology, was then quantified. zQ175 HD mice showed markedly reduced DARPP32 signal, whereas AQP4 overexpression partially preserved DARPP32 immunoreactivity (**Fig. 4E–F**).

We further determined whether AQP4 overexpression attenuated brain volume loss in zQ175 HD mice. High-resolution structural MRI was performed in 10-month-old mice. zQ175 mice exhibited significant volume loss in both the striatum (**Fig. 4G**) and whole brain (**Fig. 4H**). AQP4 overexpression significantly attenuated striatal atrophy (**Fig. 4G**), whereas whole-brain volume was not significantly restored (**Fig. 4H**).

### AQP4 overexpression improves motor coordination and attenuates body weight loss in HD mice

To determine whether the cellular and structural preservation associated with AQP4 overexpression translated into functional benefits, we assessed motor function using the balance beam test in 12-month-old zQ175 HD mice. Female zQ175 HD control mice exhibited impaired motor performance, indicated by prolonged beam-traverse time compared with sex-matched WT controls (**Fig. 5A**). AQP4 overexpression substantially reduced beam-traverse time in female zQ175 HD mice, indicating improved motor coordination (**Fig. 5A**). Male zQ175 HD mice showed a trend toward prolonged beam-traverse time compared with male WT controls, while AQP4 overexpression produced a modest improvement; however, the differences among male groups did not reach statistical significance (**Fig. 5B**).

**Figure 5.**
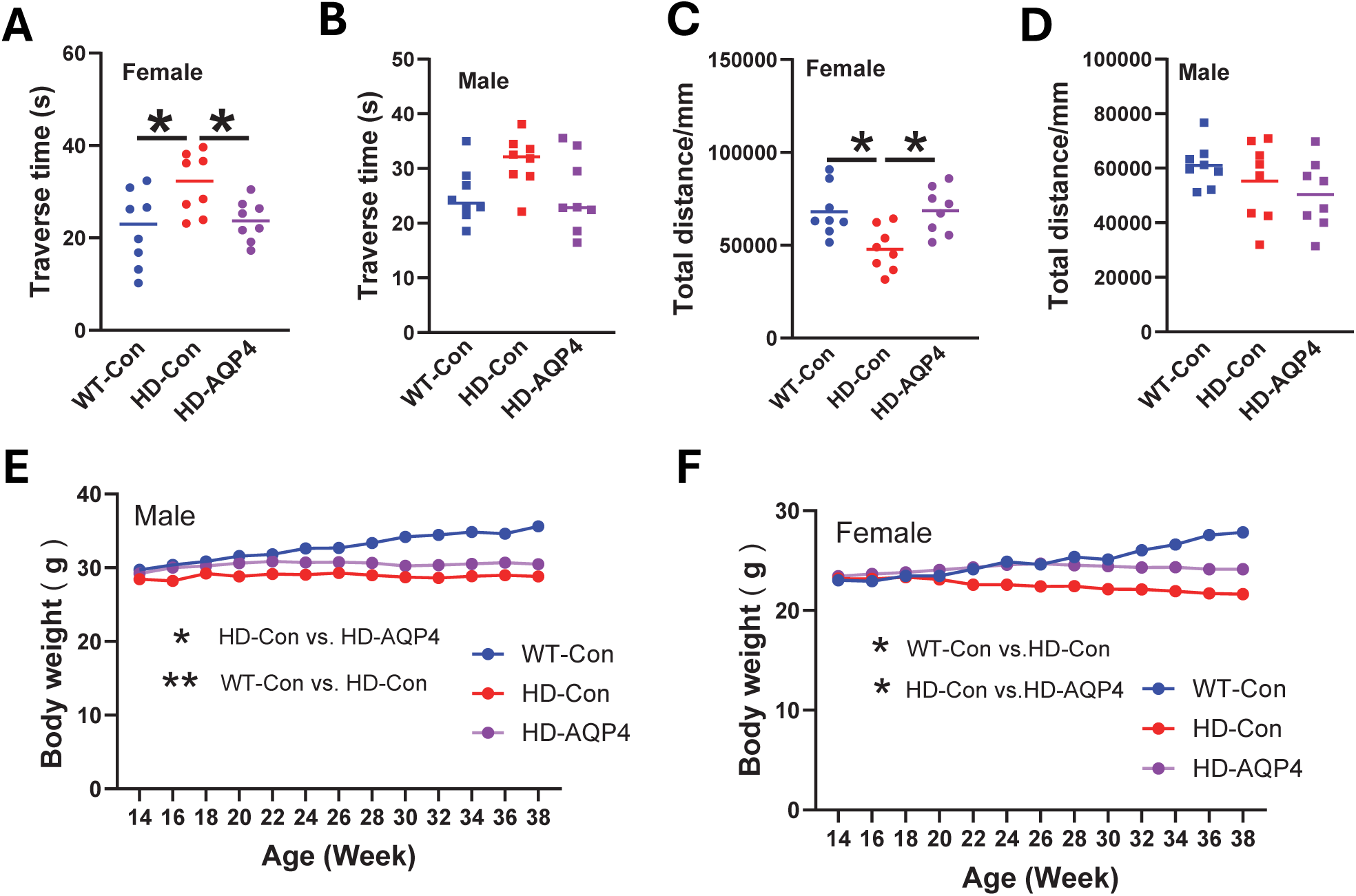
AQP4 overexpression improves motor coordination and attenuates body weight deficits in zQ175 mice. (A, **B)** Balance beam performance in female **(A)** and male **(B)** mice. Traverse time was used as a measure of motor coordination. **(C, D)** Open-field locomotor activity in female **(C)** and male **(D)** mice, measured as total distance traveled. **(E, F)** Longitudinal body weight changes in male **(E)** and female **(F)** mice. Individual data points are shown. Cross-sectional data were analyzed by one-way ANOVA followed by Bonferroni’s test. Longitudinal body weight data were analyzed using two-way repeated-measures ANOVA. *P < 0.05; **P < 0.01.

We next evaluated spontaneous locomotor activity in the open-field test. Female zQ175 HD mice exhibited reduced total travel distance compared with female WT controls, whereas AQP4 overexpression restored locomotor activity in HD mice, bringing performance toward levels observed in WT mice (**Fig. 5C**). Male zQ175 HD mice showed a modest reduction in travel distance relative to WT controls, but this difference was less pronounced. AQP4 overexpression had no significant effect on total travel distance in male zQ175 HD mice (**Fig. 5D**). Together, these behavioral findings indicate that AQP4 overexpression confers functional benefit, this effect is more pronounced in female than in male zQ175 HD mice.

Because loss of body weight maintenance is a characteristic feature of the zQ175 HD model and can emerge before overt motor impairment, we monitored body weight longitudinally. Both male and female zQ175 mice showed loss of body weight gain beginning around 22–24 weeks of age, followed by progressive divergence from sex-matched WT mice (**Fig. 5E-F**). AQP4 overexpression rescued these body weight deficits in both male and female zQ175 HD mice, with the effect being more pronounced in females.

### AQP4 overexpression has no detectable adverse effects in wild-type mice

To evaluate the potential adverse effects of sustained AQP4 overexpression in the brain, we conducted a parallel study in wild-type mice receiving AAV-AQP4. AAV-AQP4 was delivered to WT mice at 2 months of age. Longitudinal monitoring revealed no detectable effect of AAV-AQP4 administration on body weight in WT mice (**Fig. 6A–B**). We also assessed brain volumes and motor behavior in 10-month-old WT mice, corresponding to 8 months after AAV-AQP4 delivery. Structural MRI showed no significant differences in striatal or whole-brain volume (**Fig. 6C–D**). Furthermore, behavioral assessments, including the balance beam and open-field tests, revealed no detectable deficits in motor coordination or locomotor activity after AAV-AQP4 administration (**Fig. 6E–H**). Collectively, these findings indicate that sustained AQP4 overexpression was well tolerated in wild-type mice under the conditions tested.

**Figure 6.**
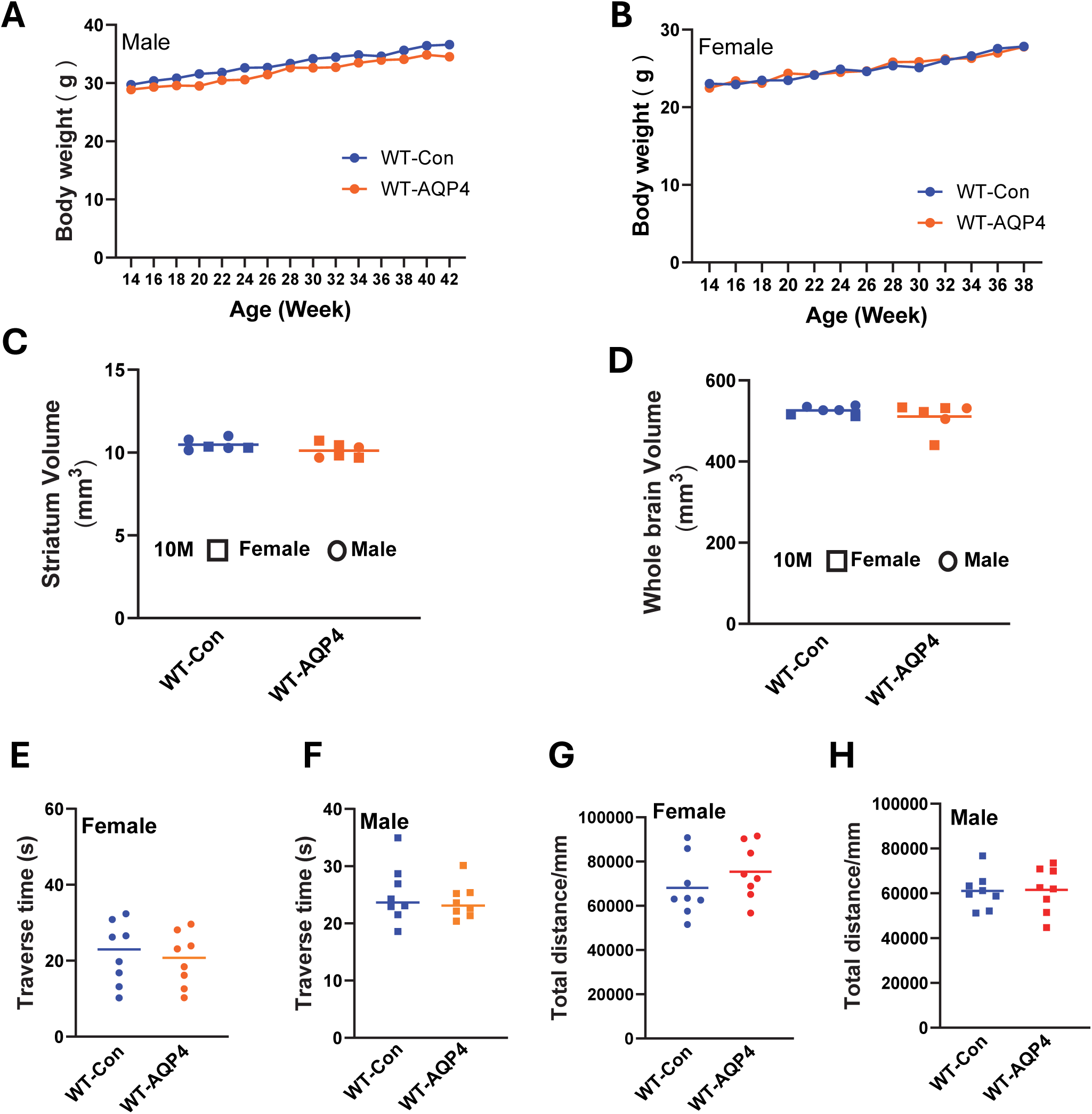
AQP4 overexpression does not produce detectable physiological, structural, or behavioral alterations in wild-type mice. **(A, B)** Longitudinal body weight measurements in male **(A)** and female **(B)** WT-Con and WT-AQP4 mice. **(C, D)** MRI-based quantification of striatal volume **(C)** and whole-brain volume **(D)**. **(E, F)** Balance beam performance in female **(E)** and male **(F)** mice. **(G, H)** Open-field locomotor activity in female **(G)** and male **(H)** mice. Individual data points are shown. Cross-sectional data were analyzed by unpaired two-tailed Student’s t-tests. Longitudinal body weight data were analyzed using two-way repeated-measures ANOVA.

## Discussion

The present study provides proof-of-concept evidence that augmenting AQP4-mediated glymphatic transport mitigates HD-associated neuropathology and behavioral deficits in a preclinical HD mouse model. We demonstrated that systemic delivery of BBB-penetrant AAV-AQP4 boosted its expression and restored its perivascular localization in zQ175 HD mice. Restored perivascular AQP4 localization results in improved glymphatic influx and downstream drainage to the dCLNs. These changes were accompanied by broad attenuation of HD-associated phenotypes, including reduced mHTT aggregate burden, lower CSF neuronal injury markers, attenuated gliosis, preservation of striatal DARPP32 immunoreactivity, and attenuated striatal atrophy. These rescue effects in neuropathology were further translated into improved motor function and body weight maintenance in HD mice without detectable adverse effects in WT mice, implicating a potential therapeutic impact of restoring glymphatic function in HD.

Efficient glymphatic transport depends on the polarized localization of AQP4 channels at vascular-facing astrocytic endfeet, where they facilitate convective CSF-ISF exchange and waste clearance ^28^. Genetic deletion or disruption of AQP4 polarization compromises parenchymal solute clearance, supporting AQP4 as an important regulator of brain fluid homeostasis ^28,29^. We previously demonstrated that glymphatic flow is perturbed in premanifest zQ175 mice, a deficit tightly correlated with the loss of perivascular AQP4 polarization ^15^. In this study, we established that exogenous AQP4 expression can restore perivascular AQP4 localization in zQ175 mice, improve glymphatic flow, and attenuate HD pathology. These data suggest that impaired glymphatic–lymphatic transport is a modifiable component of HD-associated pathology.

A key finding of this study is that AQP4 overexpression was associated with reduced striatal mHTT aggregate burden. Although mHTT is predominantly intracellular, previous studies indicate that mHTT species can be secreted into the extracellular space through unconventional endosomal/lysosomal pathways and can be cleared from the brain through active mechanisms^13,14,30^. Once extracellular, these proteotoxic species could be accessible to perivascular glymphatic clearance pathways. Our functional tracer assays showed that AQP4 overexpression improved glymphatic influx and downstream drainage. These changes may facilitate the removal of extracellular mHTT species or other neurotoxic solutes that contribute to proteostatic stress. However, because extracellular mHTT flux was not directly measured, the contribution of glymphatic–lymphatic transport to reduced mHTT aggregation remains to be determined. The parallel reductions in CSF NfL and total Tau further suggest attenuation of neuroaxonal injury. Because NfL and Tau reflect neuronal and axonal injury rather than a single disease mechanism, their reduction should be interpreted as a downstream marker of attenuated disease-associated pathology rather than direct evidence of enhanced protein clearance. Nevertheless, intracellular protein quality-control pathways, including autophagy and the ubiquitin-proteasome system, also contribute to mHTT proteostasis ^31,32^.

AQP4 overexpression was also associated with reduced striatal neuroinflammatory pathology, including tempering microgliosis and astrogliosis. Chronic exposure to localized proteotoxic stress drives astrocytic and microglial activation, generating a self-perpetuating cycle of neuroinflammation and metabolic excitotoxicity. By enhancing glymphatic–lymphatic transport, AQP4 overexpression may facilitate the removal of neurotoxic metabolites and metabolic debris, thereby attenuating astrogliosis and microgliosis.

Enhanced glymphatic influx is expected to facilitate downstream meningeal lymphatic drainage and the clearance of extracellular solutes into dCLNs, thereby improving overall brain fluid homeostasis ^33–35^. From a translational perspective, these findings introduce a non-cell-autonomous disease-modifying paradigm that complements current HTT-lowering modalities. While antisense oligonucleotides (ASOs), RNA interference, and CRISPR-based genome editing are designed to target the root genetic defect ^36–43^, they do not directly target impaired extracellular fluid transport or clearance-associated pathways. AQP4-mediated enhancement of brain fluid transport may complement HTT-lowering strategies by targeting impaired clearance and extracellular fluid homeostasis, rather than directly modifying HTT expression. Recent neuroimaging studies have suggested glymphatic deficits and altered CSF dynamics in manifest HD patients ^16–18^, suggesting that this fluid-dynamic pathway represents a clinically relevant target with translational validity. Furthermore, the lack of detectable changes in measured physiological, structural, or behavioral outcomes in WT mice suggests that overexpressing AQP4 is well tolerated.

Despite these promising outcomes, several limitations require consideration. First, although AQP4 overexpression restored glymphatic transport and reduced mHTT aggregates, our data do not definitively parse the precise contributions of intracellular versus extracellular clearance networks. Future studies deploying real-time microdialysis or advanced biosensors to track extracellular mHTT flux will be required to define these specific clearance kinetics. Because mHTT turnover is regulated by multiple intracellular protein quality-control pathways, including autophagy and the ubiquitin–proteasome system^31,32^, the contribution of glymphatic transport to mHTT proteostasis remains to be explored. Second, AQP4 is a highly multifunctional channel that modulates BBB integrity, neurovascular coupling, and extracellular potassium homeostasis ^28^. We have demonstrated impaired BBB integrity in zQ175 mice ^44^, suggesting that restoration of BBB function may also contribute to the observed neuroprotective effects. Third, the therapeutic intervention was initiated during the premanifest stage. Whether the glymphatic enhancement mediated by AQP4 overexpression can attenuate or reverse HD pathology when delivered post-symptom onset is a critical next step for defining the clinical therapeutic window. Finally, although the zQ175 knock-in model recapitulates important aspects of human HD genetics and pathology, validation across additional HD models is needed. In addition, AAV-PHP.eB-mediated CNS transduction is highly dependent on mouse genetic background and is not directly translatable to humans. Therefore, future studies should include clinically relevant delivery strategies to determine whether AQP4-based modulation of glymphatic transport can be adapted for therapeutic use ^45,46^.

In conclusion, our findings demonstrate that AQP4 overexpression enhances glymphatic dynamics, alleviates neuropathology, and mitigates motor deficits in an HD mouse model. By preserving the efficiency of perivascular waste clearance, our study provides proof-of-concept that enhancing brain glymphatic transport mitigates proteotoxic stress in HD. These findings support restoration of glymphatic homeostasis as a potential therapeutic strategy for HD.

## Data availability

The datasets generated and analyzed during the current study are available from the corresponding author upon reasonable request.

## Acknowledgements

We thank Dr. Xinping Huang at the Emory University Viral Vector Core for AAV packaging and production. We also thank Lida Du, Qian Wu, Yuan Zhou, Zhenyu Wang, and Chelsea Guo, members of Dr. Duan’s laboratory, for helpful discussions and technical assistance.

